# Potential for real-time health and welfare monitoring in experimental rabies infection in red fox (Vulpes vulpes) using implants

**DOI:** 10.64898/2026.01.16.699951

**Authors:** Julie Teresa Shapiro, Eric Morignat, Denise Dubois, Nicolas Penel, Aurélien Madouasse, Emmanuelle Robardet, Jean-Philippe Amat, Viviane Henaux, Sandrine Lesellier

## Abstract

Monitoring and improving animal health and welfare is a pressing need. Technological advances are increasing our ability to do so in a range of settings but currently, there are limited uses in experimental settings for less common model animals. Red foxes (*Vulpes vulpes*) are maintained by the French National Agency for Food, Environmental, and Occupational Health and Safety (ANSES) for regulatory and research purposes, primarily related to its role as a European Union and French Reference Laboratory for rabies and Echinococcus spp. In 2022, four foxes were surgically fitted with internal implants that recorded body temperature and activity levels and subsequently inoculated with rabies virus (RABV). Retrospectively, we tested the potential of these two variables to provide an early warning for the onset of rabies symptoms. We applied two anomaly detection algorithms (Shewhart and EWMA) with varying confidence levels to the data sets and compared how early the different models could detect significant changes while limiting false alarms during the calibration period. We hypothesized that body temperatures would rise significantly and foxes would significantly alter their activity levels at the beginning of infection, both at an earlier stage than is detectable through direct observation. We found that foxes significantly changed their activity during infection. We were best able to detect these changes using the EWMA algorithm, in some cases producing consecutive alarms up to two weeks before the death of an animal, while limiting false alarms. We found no evidence of fever in any of the infected foxes and body temperature did not appear to be a reliable indicator of foxes’ health. While here we applied our methods to a particularly severe and rarely implemented model with a very small sample size, this proof of concept illustrates the potential of these methods for a wide range of other situations that would benefit from similar long-term monitoring, including experimental protocols with milder clinical signs and routine monitoring for unexpected declines in health or welfare.

## Introduction

The analysis of behavior and physiology is a reliable and important way to assess both the health and welfare of animals. When an animal is ill, regardless of the specific pathogen, the immunological response and release of cytokines trigger a suite of common physiological and behavioral changes, including fever, a reduction in activity, and decreased appetite (Hart and Hart, 2019; Tizard, 2008). These changes are adaptive and allow sick individuals to conserve their resources and energy to fight the infection (Hart and Hart, 2019). These responses are not limited to infectious disease but may appear as a reaction to stress or impaired welfare (Chapa et al., 2020; Hart and Hart, 2019).

Sensors are a tool that can continuously and automatically collect vast amounts of data related to animal behavior and physiology. They can take a variety of forms, such as surgically implanted internal devices, ear tags, collars, and vests (Chapa et al., 2020; van Dixhoorn et al., 2024). Combined with the increased capacity to store and analyze “Big Data”, the use of sensors has greatly increased our ability to monitor individual animals in real-time. This is important because in addition to the fact that it is impossible to continuously monitor each individual, especially in large herds, without some level of automation, animals will often instinctively mask symptoms, resulting in a later detection of the issue, often when it has advanced to a more serious stage (Tizard, 2008; Weary, 2008). A variety of sensors have been developed extensively for use in farm animals as a part of precision livestock farming, particularly for cattle, for which there are already a number of commercial products (Chapa et al., 2020; van Dixhoorn et al., 2024). They may be able to monitor both physiological parameters, especially body temperature, or behaviors, including activity or basic movement, or more specific behaviors such as feeding or lying, although the latter requires more extensive validation (Chapa et al., 2020).

These tools have been applied to the early detection of diseases, particularly in livestock species. In calves, Wottlin et al., 2021 found that monitoring step counts, frequency and duration of lying, and a calculated motion index via accelerometers attached to the leg allowed detection of bovine respiratory disease up to two days earlier than manual observation. Similarly, time spent inactive, active, or lying, measured with an ear-tag accelerometer, was able to predict neonatal calf diarrhea in individuals the next day (Goharshahi et al., 2021). In an experimental group of pigs, sensors were able to detect African Swine Fever earlier than observers by signalling a sudden increase in body temperature along with decreased movement (Martínez-Avilés et al., 2017).

Real-time monitoring of animals is also essential for animals in experimental settings, who are protected by legislation (Codecasa et al., 2021), such as Directive 2010/63/EU in the European Union (European Parliament, 2010) or the Institutional Animal Care and Use Committee in the USA (US National Research Council, 2011). Sensors measuring physiological parameters, such as body temperature and heart rate in particular, are widely used for laboratory animals (Hawkins, 2014; Morton et al., 2003). The use of these technologies can contribute to the three R’s (refine, reduction, replacement) of laboratory animal use (Russell and Burch, 1959), particularly in terms of refinement (Hawkins, 2014; Knight, 2011; Morton et al., 2003; Stephens et al., 2002). Such data can be helpful in determining humane endpoints (US National Research Council, 2009) for experiments with a terminal outcome, especially if other symptoms are difficult to detect, could become particularly distressing to the animal, or appear only shortly before death (Dawson et al., 2017). They can be used to monitor animals’ welfare, stress levels, and recovery from procedures and protocols (Hawkins, 2014; Van Loo et al., 2007). Finally, they can increase the amount and quality of data collected from each individual while decreasing handling, restraint, and stress (Morton et al., 2003).

Effective real-time monitoring for both the health and welfare requires first establishing a baseline for the parameter being measured during a calibration period, which is assumed to be free of specific events that could alter the measured parameter. Once this baseline has been established, the extent of “normal” variation can be measured statistically. For data beyond this reference window, an algorithm for the detection of anomalies can be applied to determine when the parameter being measured has moved beyond the threshold of “normal” variation. Different algorithms can detect different types of changes. For example, the widely used Shewhart algorithm, first developed for quality control, measures deviations from the baseline mean. It is most useful for detecting large, abrupt changes from the mean as there is no influence of previous observations on the current observation (Nazir et al., 2013; Shewhart, 1923). Other algorithms, particularly Exponential Weighted Moving Average (EWMA) or cumulative sum (CUSUM), that incorporate past information, are better adapted to detecting more gradual but permanent shifts (Lucas and Saccucci, 1990; Nazir et al., 2013; Roberts, 1959).

While they are generally an uncommon experimental model, red foxes (*Vulpes vulpes*) are maintained by the French National Agency for Food, Environmental, and Occupational Health and Safety (ANSES) for regulatory and research purposes, primarily related to rabies and *Echinococcus* spp. Using automated, real-time monitoring systems is especially important for wild species, as they modify their behavior and mask disease symptoms even more than domesticated species in the presence of humans, which makes accurately monitoring their health and welfare through direct observation challenging, if not impossible (Carpenter, 2010). Real-time monitoring of the animals’ behavior and physiology can also detect unexpected illness or health problems not related to experimental studies.

Rabies is a zoonotic disease caused by the classical rabies virus (RABV; Genus: *Lyssavirus*, family: Rhabdoviridae). It causes acute encephalitis and is nearly always fatal in domestic and wild mammals, as well as humans once symptoms have appeared (Walker et al., 2018), although there are documented cases of asymptomatic infection in both dogs (Fekadu, 1975; Fekadu et al., 1982) and hyenas (East et al., 2001). Foxes are the primary terrestrial wildlife reservoirs of the virus in France and Europe (Freuling et al., 2013; Riccardi et al., 2021; Toma and Andral, 1977). In its role as the European Union Reference Laboratory (EURL) for Rabies, ANSES has conducted experimental inoculations of captive foxes with RABV for reference virus production that is necessary for diagnostic performance evaluation. Monitoring these foxes is particularly challenging because the asymptomatic period for rabies can extend for weeks or even months and the onset of symptoms after inoculation (or infection) is difficult to predict precisely, including at high doses (Aubert et al., 1991; Black and Lawson, 1970; Blancou et al., 1979b; George et al., 1980; Sykes-Andral, 1982; Toma and Andral, 1977). Further, the symptoms of the disease may vary widely between individuals, especially between the paralytic and furious forms. The most common symptoms include loss of appetite, lethargy, vocalizations, motor problems, nervousness, and aggressiveness, while in severe furious forms, foxes may bite objects to the point of breaking their teeth or jaws. However, any symptoms are often evident only at the end of disease development and in some cases only a few days or less before death (Blancou et al., 1979b; George et al., 1980). Furthermore, rabies experimental infections are conducted at level 3 containment, which limits the capacity of staff to monitor animals continuously. Due to these various factors, clinical signs can therefore be mistakenly reported as absent or delayed, with major impacts for the animals’ health and welfare. In light of these various challenges, real-time monitoring of these foxes using tools such as sensors and implants can help establish experimental endpoints as early and precisely as possible.

The use of implants can also support the collection of data that are difficult or dangerous to obtain otherwise due to the infectious agent or substance being studied (Dawson et al., 2017). For example, it is impractical and potentially dangerous to take body temperature measurements during experimental protocols related to rabies, particularly once symptoms have begun. References for rectal temperature in foxes are not available after experimental infection, and scarce in naive foxes (Kreeger 19888, Oppermann and Bakken, 1997). In addition, the activity kinetics of experimentally inoculated foxes has never been quantified, only described qualitatively.

In 2022, the EURL for Rabies conducted a study that required the inoculation of four foxes with rabies virus (RABV). They were all fitted with internal implants that measured body temperature and activity. While these parameters were measured in real-time, we were only able to analyze them retrospectively. Our objectives were: 1. To describe the body temperature and activity of the foxes during the development of rabies, which to our knowledge has not previously been published and 2. To test the potential of both body temperature and activity data to provide an early warning for the onset of rabies symptoms by applying two anomaly detection algorithms (Shewhart and EWMA) to the data sets. We compared the potential of each variable and algorithm to provide an early detection of significant changes during the monitoring period while limiting false alarms during the calibration period. We hypothesized that body temperatures would rise significantly and that foxes would significantly alter their activity, both at an earlier stage than is detectable through direct observation. Using this study as a proof of concept, we illustrate how the collection of such data and their analysis could be leveraged for real-time monitoring capable of providing an early warning for the onset of disease symptoms before they are detected by observation alone.

## Methods

### Experimental set-up

The experimental protocols in foxes were authorized under European legislation by the French Ministry of Research APAFIS #34638-2021122119167519. A total of four foxes were included in the study with the following identifiers : 0fae, 0faf, 0fb0, and 0fb1.

On 4 July 2022, the four foxes were fitted with easyTEL+ implants (model ET-M2-ETA - Emka Technologies, France) (see Supplementary Figure S1) (Shapiro et al., 2026) while under general anesthesia with intramuscular medetomidine chlorhydrate (0,1mg/kg) and ketamine chlorhydrate (10mg/kg) followed by isoflurane maintenance. Each implant was composed of a silicone outer shell, weighing 17.4 g (dimensions 26.90 x 15.80 x 22), with two 30 cm-1mm electrodes and a battery allowing a maximum of 175 days autonomy. The implant was programmed to monitor and record body acceleration, temperature (°C), and electrocardiogram (ECG) with continuous radio-telemetry transmission of raw signals. A telemetry receiver was positioned outside the cages in the range of 3-5m from the animal carrying the implant. The data collected by the transmitter were transferred to a computer in a protected format from which values averaged over 10 seconds were exported to a text file (.txt). Derived parameters (such as heart rate) or averaged values were automatically saved every 10s into datafiles.Temperature was measured with a range of 4°-50°C with a resolution of 0.04°C. Body acceleration was measured by the implant using an embedded triaxial accelerometer which measures acceleration in three axes (x: side to side acceleration; y: up and down acceleration; z: front and back acceleration). Any movement of the implant results in changes to the acceleration in each axis. Total body acceleration was calculated by Emka Technologies’ IOX2 acquisition software by calculating the modulus of each axis’ acceleration 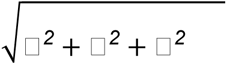 (*g*s, e.g. gravity acceleration, m/s^2^) (Emka Technologies, 2025; Schumacher et al., 2019).

Each implant was positioned in a muscular pouch between the abdominal rectus muscle and transversus abdominis muscles. The ECG electrodes were placed on the thorax in a modified led II configuration. The electrodes were exteriorized and the pouch was closed with synthetic absorbable suture poliglecaprone 25 (Monocryl 4/0, Ethicon J&J MedTech, USA). The negative electrode was tunnelized under the skin from the muscular pouch to the upper right thoracic area. A small boucle of electrode was prepared, placed under a small pocket of thoracic muscles, and secured in place with Prolene 5/0 (Ethicon J&J MedTech, USA). The positive lead was tunnelized under the skin and guided approximately 5 cm left from the xyphoid process. A small loop of electrode was prepared, placed under a small pocket of muscles and secured in place with 5/0 Prolene. The subcutaneous tissue and the skin of all implantation sites were closed layer by layer with Monocryl 4/0 (continuous pattern, intracutaneous for the skin). Each fox was treated with tolfenamic acid (1mL/10kg) and amoxicillin (1mL/kg) for the three days following implantation to reduce local infection and inflammation.

On 8 July 2022, each fox was inoculated intramuscularly with classical rabies virus (RABV) batch Salivary Gland (SG) 5-4, as described in (Aubert et al., 1991). This RABV batch had been isolated in France in 1982 from a naturally infected fox and then produced in-house through passage in a fox host, the salivary glands of which were homogenised, encapsulated in a sealed glass container, and stored in liquid nitrogen from 18/10/1982 (Aubert et al., 1991). Before inoculation for this study, the viral stock was thawed in a dark environment and diluted to 103.9 DL 50 (Intra Cerebral inoculated *Souris* (mouse) (ICS)). The viral suspension was injected IM in the left temporal muscle of each anesthetised fox with medetomidine chlorhydrate (0,1mg/kg) and ketamine chlorhydrate (10mg/kg). Anesthesia was reversed by intramuscular atipamezole (1mg/kg).

After inoculation, all animals were closely monitored by animal caretakers at least once a day before clinical signs were reported, and at least twice a day after the development of clinical signs. The animals were observed and any clinical symptoms of rabies, such as anorexia, paralysis, excitability, nervousness, or biting, were noted as present/absent on paper scoring sheets as described in George et al., 1980.

As this was the first deployment of the implants for an experimental protocol of rabies infection in foxes and the incubation period may extend for several months (Aubert et al., 1991; Black and Lawson, 1970; Blancou et al., 1979b; George et al., 1980; Toma and Andral, 1977), there were concerns regarding battery life of the implants. For this reason, the implants were turned off for twelve days starting on July 27th and turned back on on August 9. No data was collected by the implants for any animals for twelve days from 28 July - 8 August, inclusive. Because endpoints were defined based on caretakers’ observations, this gap in data collection from the implants did not impact the animals’ health or welfare. Animals remained under observation by caretakers as planned under ethical authorisation for any symptoms during the entire study period.

Three foxes (0fb1, 0faf and 0fae) developed typical rabies symptoms that were noted by animal caretakers on 3, 5 and 17 August 2022, respectively (Table 1). According to the planned protocol, 0fae received acepromazine 1mg/kg and buprenorphine (0.1mg/kg) on the day of euthanasia (18 August 2022) to alleviate detected symptoms associated with incoordination, as did 0faf two days before death (6 August 2022). 0faf and 0fae were euthanised under general anesthesia when symptoms progressed to the more severe experimental end point on 8 and 18 August 2022, respectively. 0fb1 was found to have died on 4 August 2022, one day after mild symptoms were detected by direct observations (3 August 2022). Post-mortem infection by RABV was confirmed for all three foxes with the FAT (fluorescence antibody technique) test on central nervous system tissue followed by confirmation of the RABV batch used in the inoculation (SG 5-4) via PCR (WOAH, 2023).

**Table 1.**
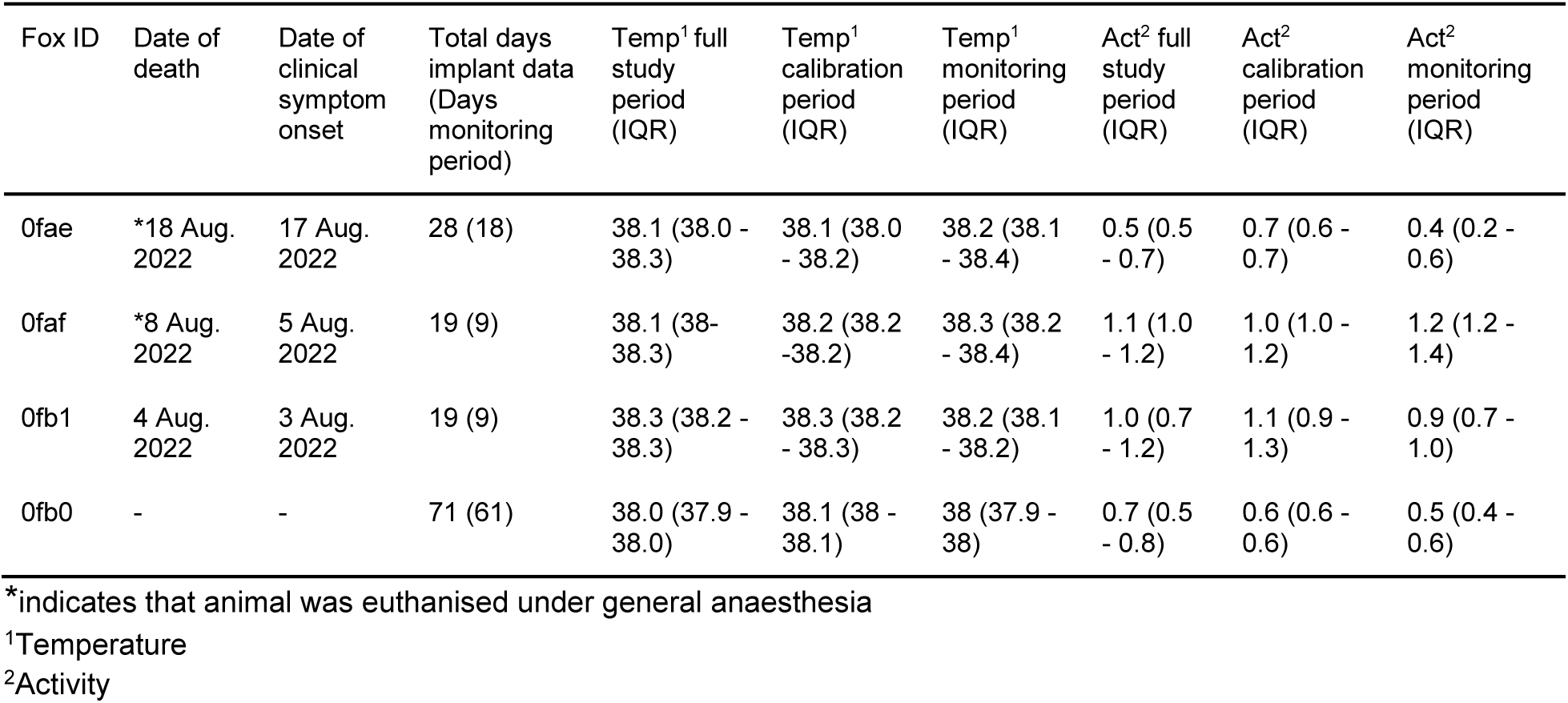
Description and summary statistics (median and inter-quartile range (IQR)) of body temperature (Temp.) and Activity (Act.) data available for the four foxes included in this study. We also note the ID, date of death, and the day clinical symptoms were observed by animal caretakers. The number of days of data does not include the initial four days that were excluded from the calibration period.

Fox 0fb0 did not develop any symptoms of rabies. This fox is still alive at the time of writing (January 2026) and has never shown any clinical signs of rabies, allowing this fox to act as a negative control.

### Statistical analyses

All data processing and analyses were conducted using the software R version 4.1.2 (R Core Team, 2013). All data and scripts are available at https://doi.org/10.5281/ZENODO.18258117 (Shapiro, 2026).

The raw data were recorded every 10 seconds. However, because this scale would introduce too much noise, we performed an initial data aggregation for both temperature and body acceleration of 15 minutes, an approach which has been used in similar studies for other species (Wottlin et al., 2021). For body temperature we calculated the mean temperature over each 15-minute interval (°C), while we used the sum of body acceleration (*g*s) for each 15-minute interval to measure activity.

For each fox, we divided the data into two time periods: a 10-day calibration period to establish a baseline starting the day after inoculation (9 - 18 July 2022), followed by a monitoring period from 19 July 2022 until each animal’s death. We excluded the days between the placement of implants to the day of inoculation (4 - 8 July 2022) because the time between the two procedures was very short and included time under anaesthesia, which introduced too much noise and variability into the data to provide a reasonable baseline. The minimal incubation period for rabies in foxes is around 12 days, although it may extend for several months (Aubert et al., 1991; Black and Lawson, 1970; Blancou et al., 1979b; George et al., 1980; Toma and Andral, 1977). Therefore we could reasonably assume that any alarms during the 10-day calibration period were not due to symptoms of rabies infection and thus false positives.

In order to characterize body temperature and activity, we calculated summary statistics of these variables for each fox over the study period. For each fox, we then compared its daily body temperature and activity level during the calibration and monitoring periods using Welch’s two-sample t-test to determine if these measures varied significantly over the two time periods.

We then used two statistical process control algorithms (Shewhart and the Exponentially Weighted Moving Average (EWMA)) to detect significant changes in temperature and activity. Both algorithms function by calculating an upper and lower limit based on the mean (Shewhart) or a weighted moving average (EWMA) during the calibration period, plus and minus a margin based on the variance of the data and the specified confidence level (Shewhart) or number of standard deviations (EWMA). We divided the data into three time periods, to which we applied both the Shewhart and EWMA algorithms: daily (24h), day-time (6h - 20h59), and night-time (21h - 5h59 the following day), in order to account for potential differences in physiology and behavior between day-time and night (Ahola et al., 2001; Díaz-Ruiz et al., 2016; Kreeger et al., 1989; Monterroso et al., 2014; Oppermann Moe and Bakken, 1997). For each time period after the calibration period, the mean (Shewhart) or weighted moving average (EWMA) of the time period is calculated from the values of each 15-min interval in the given time period. If the mean or weighted moving average is above the upper limit or below the lower limit, the value is considered to be outside of the normal range and thus triggers an alarm.

We implemented the Shewhart and EWMA algorithms using the functions “qcc” (Shewhart) and “ewma” of the package qcc (Scrucca, 2004) in R (R Core Team, 2013). For each time period (daily, day-time, night-time), we tested three confidence levels for each algorithm, for Shewhart : 0.95, 0.997, 0.9999 and their equivalents in terms of the number of standard deviations (nσ, also known as L) for EWMA: 2, 3, 4. For EWMA we also specified gamma, the smoothing parameter, which determines the rate of decay when calculating the weighted average and takes a value between 0 and 1. We set gamma = 0.3 as values between 0.25 - 0.5 are considered the most appropriate for relatively quickly detecting shifts of at least nσ=2, typical of most applications of statistical process control (Lucas and Saccucci, 1990). Higher values of gamma (i.e. closer to 1) approximate the Shewhart algorithm and would thus be redundant, while lower values (i.e. closer to 0) are better for quickly detecting smaller shifts around nσ=1 and would likely not be specific enough and lead to false alarms; with gamma = 0.3 such small shifts can still be detected if they are sustained over a longer period of time (Lucas and Saccucci, 1990). In total, thirty-six models were implemented for each fox.

As noted above, no data was collected by the implants for any animals from 28 July - 8 August 2022 (see Experimental set-up). We excluded these twelve days with missing data from all calculations as neither algorithm can incorporate a day without at least one data point. While this has no direct effect on the functioning of the Shewhart algorithm, as the mean is independently calculated for each day, for EWMA, the weighted moving averages after the data gap would incorporate the days before the gap in the calculation as if there had not been a gap.

In order to determine how potentially effective or reliable each variable (body temperature and activity), algorithm, confidence levels, and time periods might be for real-time monitoring, we compared the results obtained for all models and ranked their performance based on three criteria applied sequentially. Ideal surveillance should maximize true positives while minimizing false positives. In our context, we considered all alarms during the calibration period as false alarms while all alarms during the monitoring period were considered true alarms. Our first criteria for evaluating each model was the number of alarms during the calibration period, which ideally should be zero. All models that yielded zero alarms during the calibration period were ranked above any models with alarms during the calibration period.

Next, for alarms during the monitoring period, we considered multiple consecutive alarms, particularly preceding death, to be the most trustworthy true alarms because they indicate a sustained, continuous change in the variables being measured (body temperature or activity). Therefore, after considering the number of alarms in the calibration period, as our second criterion, we then considered the greatest number of consecutive alarms before the death of the animal. For the two animals (0faf and 0fb10) for which data were unavailable in the days before death because the implants were not collecting data during that period, we considered the number of consecutive alarms starting from the last day of collection.

As our final criterion, we considered the number of “singleton” alarms (e.g. a single alarm preceded and followed by a value in the normal range) during the monitoring period produced, which we aimed to minimize. While we did not necessarily consider these singletons to be false alarms, the immediate return to values within the normal range may indicate a lack of specificity. Thus, in the case of two models with an equally low number of alarms in the calibration period and an equally high number of consecutive alarms before death, we would then rank them based on the number of singleton alarms, with the model with less singleton alarms ranked higher.

## Results

Four foxes were fitted with implants and three of them developed typical rabies symptoms after inoculation with the rabies virus (Table 1). Three infected foxes died between 4 - 18 August 2022 and provided 19-28 days of implant data that were analyzed. The 4th fox (0fb0) that did not develop symptoms provided implant data for the longest period of time (71 days). No data were collected from any of the four foxes from 28 July - 8 August 2022. For the fox (0fae) which died at the latest date (18 Aug 2022), the collected data included the nine days preceding its death (Table 1). For the two foxes that died on 4 August and 8 August, no data could be collected for the 8-12 days before death. The fourth fox, 0fb0, survived inoculation and data was collected until 30 September 2022.

### Body temperature

Daily (24h) body temperature of foxes ranged from 36.9°C - 38.7°C over the study period: 37.9°C - 38.5°C during the calibration period and 36.9°C - 38.7°C during the monitoring period (Table 1, Figure 1). The highest recorded body temperature over a 24h period was 38.7°C (0fae). Even for 15-min intervals during the monitoring period, body temperature never reached a true fever, with maximum temperatures ranging from 39.4°C - 39.9°C. Temperatures up to 40°C have been measured in healthy foxes (Ahola et al., 2000; Kreeger et al., 1989) and maximum temperatures during the calibration period were almost as high at 39.1°C - 39.4°C.

**Figure 1.**
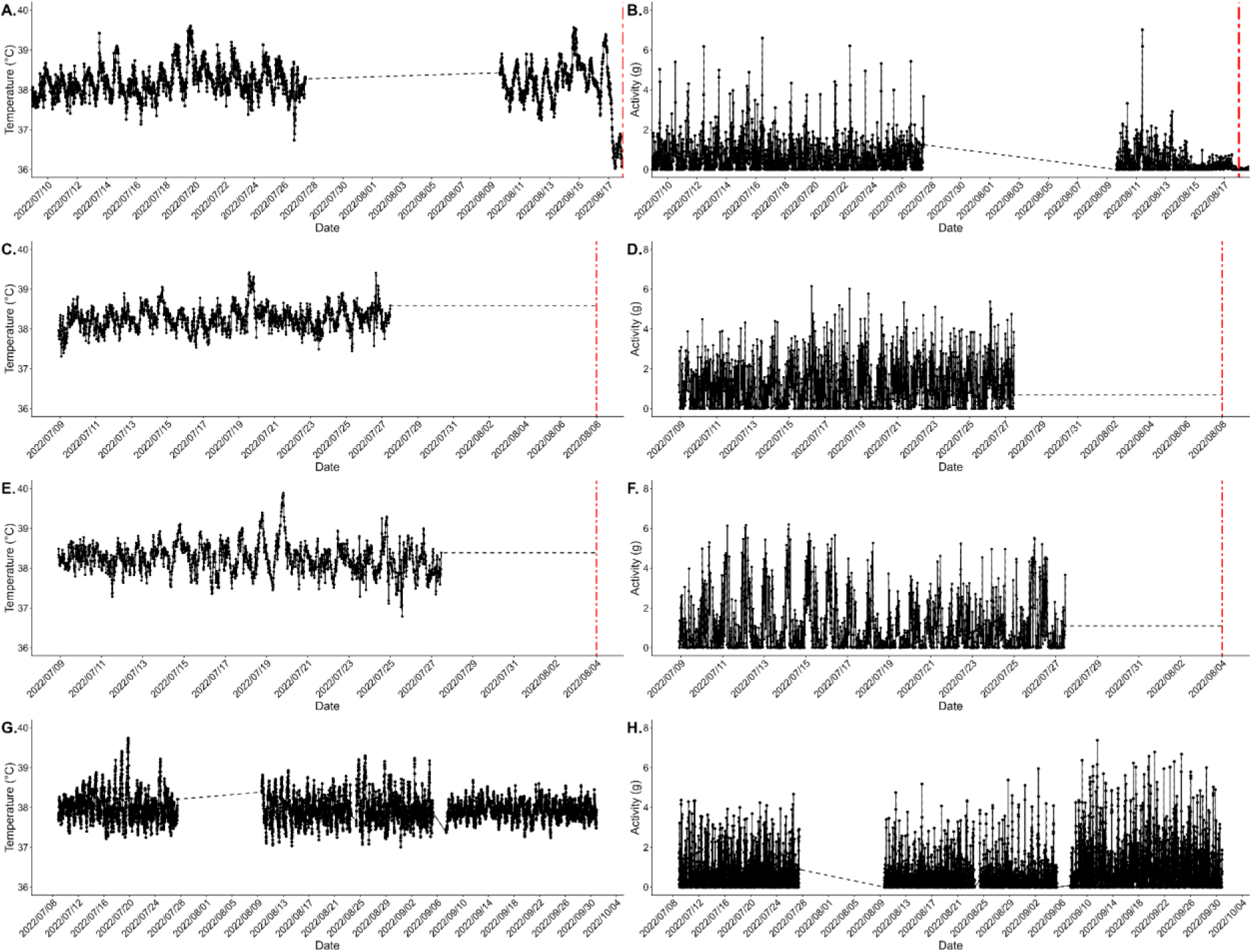
Implant data collected over the full study period for the four foxes aggregated by 15 minutes: body temperature (A., C., E., G.) and activity (B., D., F., H.) for foxes 0fae (A., B.), 0faf (C., D.), 0fb1 (E., F.), and 0fb0 (G., H.). The simple dotted black line indicates the gap in data collection from July 28 - August 8 (see Methods). The red vertical dashed-dot lines indicate the day of death for foxes 0fae (A., B.), 0faf (C., D.), and 0fb1 (E., F.). Foxes 0faf and 0fb1 died before data collection from implants resumed.

We found no significant differences in body temperature between the calibration and monitoring periods for the three foxes that died (0fae: t= −0.70, p = 0.5; 0faf: t= −1.5, p=0.14; 0fb1: t = 1.75, p=0.11). For fox 0fb0, which survived inoculation, body temperature was slightly, but significantly lower during the monitoring period (38.0°C) compared to the calibration period (38.1°C) (t=3.31, p=0.01).

### Activity

Daily (24h) mean activity levels ranged from 0.01 (on the day of death of 0fae) - 1.6 over the study period (Table 1, Figure 1). Activity was significantly higher in the calibration period compared to the monitoring period for both 0fae (t = 3.6, p = 0.001) and 0fb1 (t = 2.3, p = 0.04) while 0faf’s activity was significantly higher during the monitoring period compared to the calibration period (t = −3.5, p = 0.004). For 0fb0, the surviving fox, there was no significant difference in activity between the calibration and monitoring periods (t = −1.6, p = 0.11) (Table 1).

### Potential for real-time monitoring using Statistical Process Control

The performances of each model for anomaly detection based on the lowest number of alarms during the calibration period, the greatest number of consecutive alarms before death, and the lowest number of single alarms in the monitoring period are shown in Supplementary Tables S1-4 (Shapiro et al., 2026). Overall we found that activity was a more reliable indicator than temperature. EWMA yielded more reliable results than the Shewhart algorithm, indicating that shifts during the monitoring period were generally cumulative and sustained rather than sudden. The earliest consistent alarms appeared 9 - 16 days before clinical symptoms were observed.The best value of nσ varied from 2 - 4 between foxes. This is related to how variable the behavior was during the calibration period; a higher value of nσ decreases the number of alarms in the calibration period for individuals with more variable behavior, while maintaining a high number of consecutive alarms preceding death (Supplementary Tables S1-4) (Shapiro et al., 2026).

For fox 0fae, for which we had body temperature and activity data until death, daily (24h) activity monitoring using EWMA with nσ = 2 provided an alarm for nine consecutive days before death (10 consecutive alarms total including on the day of death). For all these alarms, activity remained below the lower limit, with a notable, continuous decrease during the last seven days as it developed the paralytic form of rabies (Figure 2). The first alarms, also related to a decrease in activity during two and three consecutive days, occurred 27 days before death, far earlier than the first clinical symptoms observed directly by animal caretakers (1 day before death). There were no alarms during the calibration period with these parameters.

**Figure 2.**
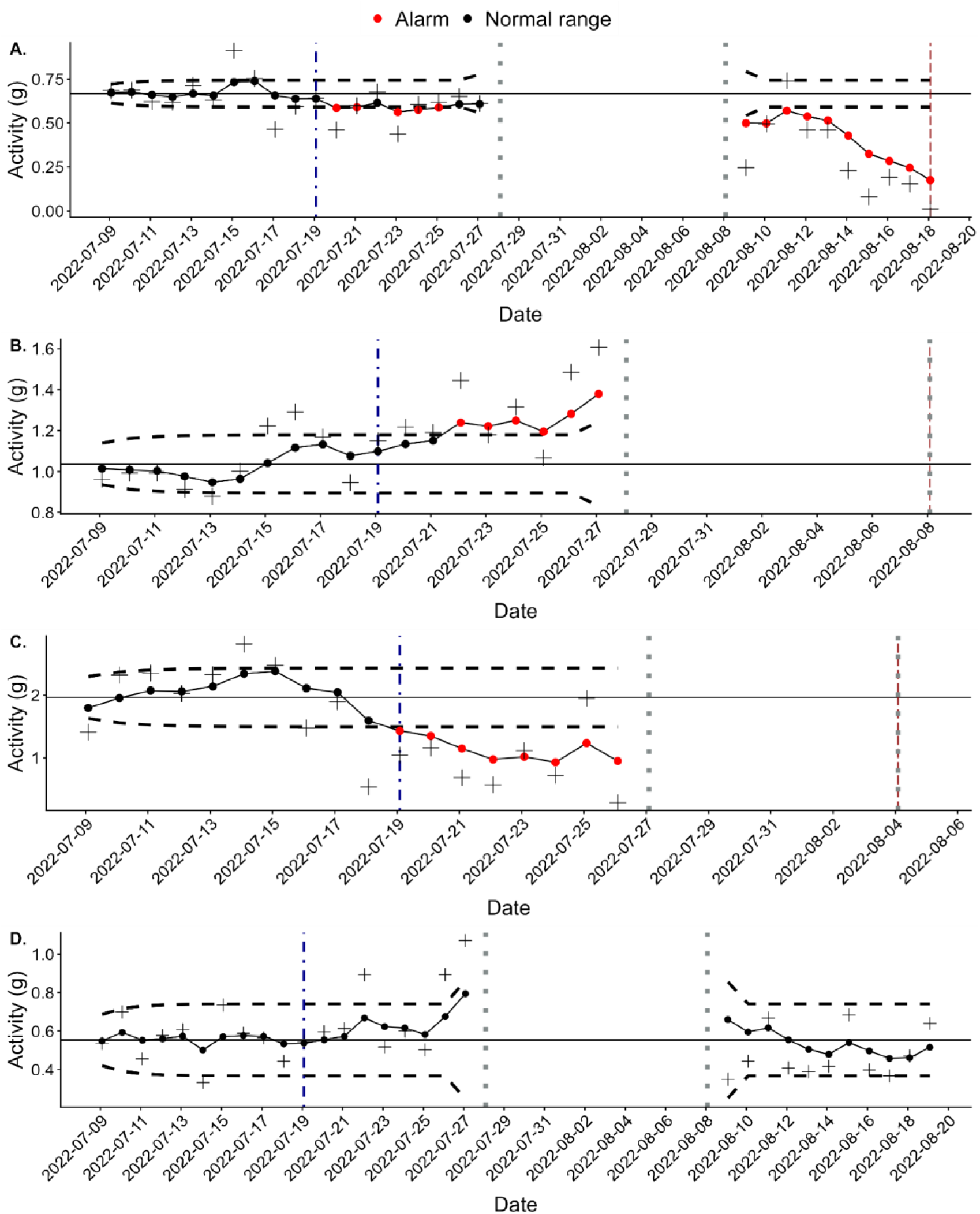
Best-performing statistical process control charts for each fox, applying the Exponential Weighted Moving Average (EWMA) algorithm to activity: A. fox 0fae (daily, nσ = 2); B. fox 0faf (daily, nσ = 3); C. fox 0fb1 (night-time, nσ = 4); D. fox 0fb0 (day-time, nσ = 4). Each dot represents the EWMA statistic for a given time unit while crosses indicate the observed value for the same time unit. The solid horizontal line represents the mean and the horizontal dashed lines represent the upper and lower limits for normal values (in-control process). Black dots indicate values in the normal range, while red dots indicate an alarm. The dark blue dot-dashed vertical line indicates the first day of the monitoring period (following the calibration period). The dark gray dotted lines indicate the days when data was not collected by the implants. The dark red dashed lines indicate the day of death for foxes 0fae, 0faf, and 0fb1 (A., B., C.). Note that due to differing levels of variability, the scale of the y-axis is different for each fox.

Similarly, for fox 0faf, the most reliable results came from monitoring daily (24h) activity with EWMA and nσ = 3, with six consecutive alarms, all above the upper limit, no alarms during the calibration period, and no singleton alarms during the monitoring period (Figure 2). The first alarm occurred 17 days before death and continued until the end of data collection twelve days before the animal’s death. These alarms correspond to elevated activity levels, consistent with the development of the furious form of rabies observed by animal caretakers. The first alarm occurred 14 days earlier than the first symptoms observed by animal caretakers (3 days before death).

For fox 0fb1, night-time activity monitored with EWMA (nσ = 4) was the best model. It provided eight consecutive alarms at the end of the data collection period, with a marked decrease in activity (Figure 2). The period with alarms corresponds to 17 days before death and 16 days before clinical symptoms were observed by caretakers (with no data during the 8 days preceding death). Again, the alarms occurred much earlier than the first symptoms identified by direct observation (1 day before death) when 0fb1 developed the furious form of rabies, although clinical symptoms remained mild. There were no alarms in the calibration period and no singleton alarms during the calibration period.

For the surviving fox (0fb0), day-time activity monitored using EWMA (nσ = 4) was the most reliable model, with no alarms during the calibration period and no singleton alarms during the monitoring period. All alarms for 0fb0 began on 9 September 2022 and continued until the end of data collection on 30 September 2022 (Figure 3). This increase in activity coincided with the introduction of new foxes and an accompanying increase in human activity related to them in the neighboring cages after a period of several weeks with no foxes in the vicinity following the deaths of the other foxes in this study (0fae, 0faf, and 0fb1). The elevated activity level during this period could possibly represent 0fbo’s new baseline after recovery from the inoculation, although the decreasing trend in the last seven days of data collection points to the more likely explanation that this was an unusually active period.

**Figure 3.**
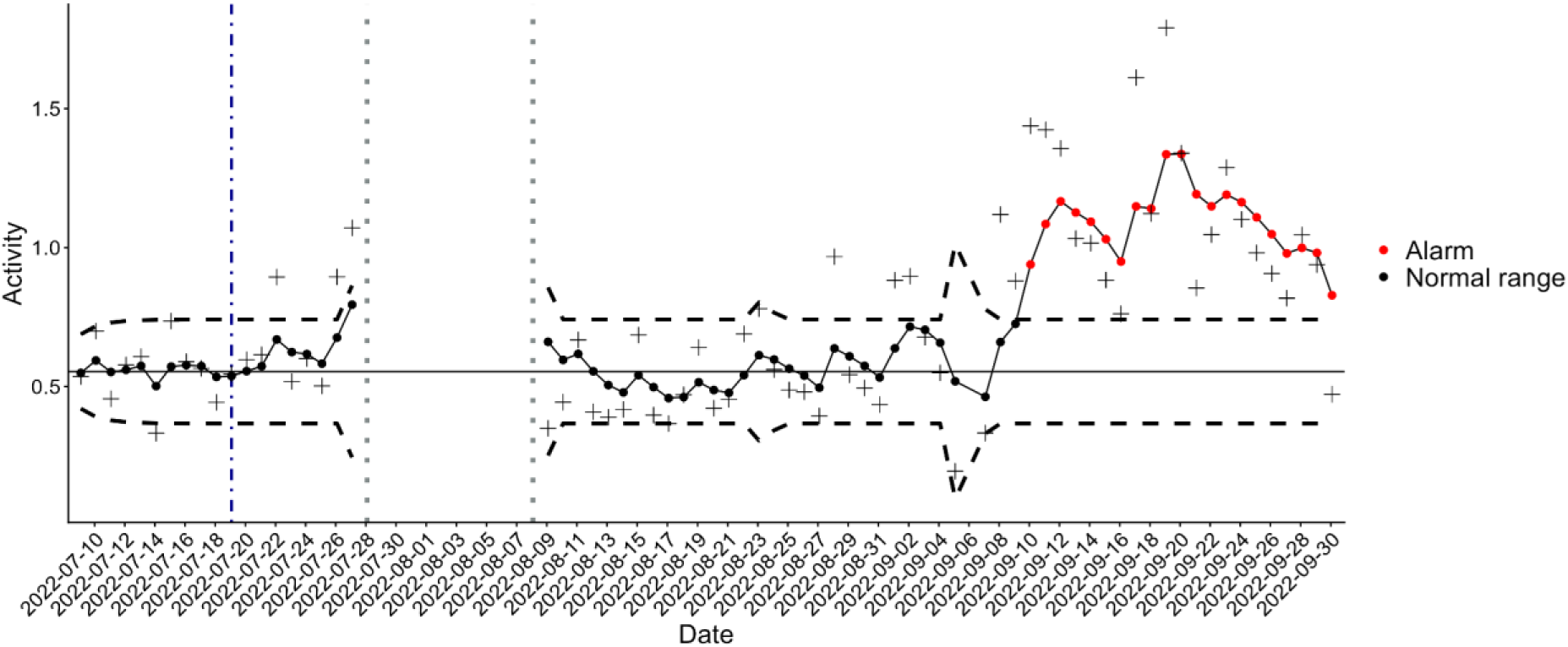
Representation of the best model results (EWMA, day-time, nσ = 4) for full data collection period for fox 0fb0. Each dot represents the EWMA statistic for a given time unit while crosses indicate the observed value for the same time unit. The solid horizontal line represents the mean and the horizontal dashed lines represent the upper and lower limits for normal values (in-control process). Black dots indicate values in the normal range, while red dots indicate an alarm. The left-most dark blue dot-dashed vertical line indicates the first day of the monitoring period (following the calibration period). The dark gray dotted lines indicate the days when data was not collected by the implant.

## Discussion

Here we have presented results from a preliminary, proof-of-concept study that demonstrates that with the right calibration of models, it is possible to produce early warnings of changes in the health or welfare of foxes in captivity and experimental settings using implants measuring physiological parameters and activity levels. Using this approach, successions of consecutive alarms were generated in three of the rabies-inoculated foxes several days or weeks (9-16 days) before clinical signs were observed by animal caretakers. This approach therefore seems to provide a potentially powerful tool to indicate the development of the infection, and that the fox may have started to experience potential discomfort. Further developing and implementing such real-time monitoring of individuals would greatly enhance the potential to monitor closely the well-being of each animal, and to accurately identify early subtle end-points triggering the gradual administration of palliative care. While further research is needed to establish the general range of precocity, this approach shows great promise to help refine experimental protocols.

While internal implants in particular should be used judiciously due to the invasiveness of the surgery needed to place them and the associated risks of infection (Hawkins, 2014; Morton et al., 2003), they may be particularly useful when inoculating high-risk pathogens (Dawson et al., 2017), such as rabies, that limits the ability to handle or interact with the animals. We used this severe and rarely implemented model to validate the system, but it could also be applicable to other situations with milder clinical signs.

We found that monitoring activity provided clearer and more consistent early warnings than body temperature for all four foxes. Our measurements of body temperature during both the calibration and monitoring periods were lower than what has been previously described in some studies (Ahola et al., 2000; Kreeger et al., 1989) but more consistent with others (Bakken et al., 1999; Oppermann Moe and Bakken, 1997), although those previous studies measured foxes that were not inoculated with any infectious agents. While we expected to see increases in body temperature as a clinical sign of inflammation and stress, studies in dogs have also shown that during rabies infection, fever, if present at all, generally remains low (Kumar et al., 2023; Nandi and Kumar, 2011). To our knowledge these are the first measures of body temperature of foxes during rabies infection.

The three foxes in this study that developed clinical signs exhibited the two typical forms of rabies symptoms - one manifesting through paralysis while the other two were marked by excitability (George et al., 1980; Sykes-Andral, 1982). Under experimental conditions, it is essential to alleviate the physical and mental stress that may be associated with symptoms, including at very early stages of the disease. However, in the case of rabies, symptoms are usually detected only a few days, or even less, before death may occur, at a well advanced stage of the disease (George et al., 1980; Sykes-Andral, 1982). Here we show that a significant change of activity can potentially be used as a quantitative, objective sign that an animal is feeling unwell and requires additional attention and treatment, even in absence of clear visible symptoms. It can also be envisaged that illness may not be the only reason for such changes, as seen in the case of 0fb0. In experimental studies when the cause of disease is clear, such a non-specific parameter is a powerful predictor of disease progression.

One of the inoculated foxes did not develop any symptoms of rabies during the course of the study or at any time afterwards and is in fact still alive at the time of writing (January 2026). While rabies is generally considered to be virtually 100% fatal, this may in fact not always be the case, particularly if symptoms do not appear. A previous inoculation study of foxes indicates that of the 50 individuals inoculated, 42 developed clinical symptoms and died (Blancou et al., 1979a). A serological study in Alaska found that Arctic foxes may be exposed to rabies and develop antibodies with no apparent symptoms and no virus found in the brain (Ballard et al., 2001). There is also evidence that dogs (Fekadu, 1975; Fekadu et al., 1982) and hyenas (East et al., 2001) can survive rabies inoculation or exposure with no symptoms, even when the virus is detected in saliva.

By comparing each animal to its own baseline and using a personalized value of nσ, we were able to tailor our models to each individual. Wottlin et al., 2021, who used several variables based on activity patterns in calves for the early detection of bovine respiratory disease, employed the same confidence level to determine the limits for all individual calves. Although their choice of confidence level was based on balancing sensitivity and specificity to maximize overall accuracy, they found that anomaly detection was less successful for individuals which displayed more variability during the baseline period because their limits were wider, leading to false negatives – a lack of alarms when those animals were in fact ill. Conversely, in our case, we found that some individuals had such high variability between days that lower values of nσ resulted in limits that were too narrow and yielded false positive alarms during the calibration period. By using a larger value of nσ and thus wider limits for these individuals, we could minimize or even eliminate the false positives while still producing early warning alarms during the monitoring period. For foxes displaying less variability, we could use a smaller value of nσ, resulting in narrower limits to preserve sensitivity without sacrificing specificity.

In addition to varying the limits, we also evaluated the potential benefits provided by the analysis of both body temperature and activity for three different time periods: complete 24-hour daily period, day-time only, or night-time only. Here again we found individual variation. For two foxes (0fae and 0faf), daily (24h) activity was the best metric, while for 0fb0 the best metric was day-time activity. While foxes in the wild are generally crepuscular (Díaz-Ruiz et al., 2016), individuals, including those in captivity (Ahola et al., 2001), may shift their activity toward either the day-time or night-time, depending on the environment, habitat, or resources available (Cavallani and Lovari, 1994; Monterroso et al., 2014). For fox 0fb1, which developed the furious form of rabies, an increase in day-time activity was observed, which was consistent with the symptoms observed by the animal caretakers. However our results also found a decrease in night-time activity (Supplementary Table S3) (Shapiro et al., 2026), which may be explained by lower levels of stimulating factors such as noise and activity at night. This result highlights the importance of the temporal period and scale in analyses. Considering multiple time periods as we have done (daily, day-time, night-time) is another way to take individual variation into account and tailor such monitoring.

Individualizing the models in this way can be a practical way to minimize both false positives and false negatives. However, in this study we clearly had the benefit of hindsight for our retrospective analyses. In an operational system, choosing the most appropriate models and parameters for each individual a priori would be challenging if not impossible. A long calibration period of at least several weeks before any manipulations or protocols are implemented can help determine individual variability, select appropriate confidence levels, and possibly adjust them if needed. Further research is needed to determine if there are certain parameters that would be useful for a significant proportion of individuals or for example if foxes might fall into different typical categories which could offer better guidance on the models and parameters to use for real-time monitoring. Here, we have only a very small sample size of four individuals, each exhibiting different levels of activity during the calibration period, in terms of both mean and variability (Table 1), as well as differing results for the most effective time period for monitoring (daily, day-time, night-time).

In this study, we analyzed two indicators, body temperature and activity, that are not specific to rabies. This study can be considered an application of syndromic surveillance, in which non-specific indicators are analyzed to detect anomalies in the health or welfare of populations (Henning, 2004), to individuals. In syndromic surveillance in general, an alarm indicates a change in a baseline but not the actual cause, only an investigation can elucidate the driver; the broader the indicator, the greater the range of probable causes (Katz et al., 2011; Mandl et al., 2004; Shapiro et al., 2025). This is also the case for individual-based monitoring, which should be used as a support tool for animal caretakers, in no way replacing their knowledge and expertise. During an experimental protocol when symptoms are expected, we can be reasonably sure of the cause and take more targeted, specific actions. But in some cases the cause of an alarm is unclear, for example outside the typical symptomatic period for a given pathogen. Such an alarm may indicate the development of a health or welfare problem not related to the experimental protocol. Finding the cause in these cases often requires a more thorough investigation or broader action, such as a clinical examination, a blood test or more frequent visual surveillance. This is illustrated by the results for fox 0fb0, which did not develop any clinical signs of rabies and had no alarms related to activity during the expected symptomatic period but exhibited increased activity for several weeks during the post-symptomatic period. We believe this increase in activity was associated with the introduction of foxes in the surrounding cages or possibly the increased human activity around it. This shows the potential capacity of the system to detect changes with different underlying causes beyond those associated with a specific protocol, including changes of state or reactions to the environment that are not necessarily negative or problematic.

Our study has a number of limitations, the most substantial being a too-short baseline period. Ideally there would have been a much longer time, at least several weeks, to collect data from the implants and establish the baseline for each fox before the rabies inoculation, which was not possible in this preliminary study. Further research is needed to determine the appropriate minimal time needed for the calibration period to establish the most normal baseline possible for both body temperature and activity. While rabies inoculation at the facility usually involves a small number of foxes, further research with other infectious agents including more individuals could further validate and refine the system, including determining if particular indicators, time periods, or model parameters are more likely to function well for the largest number of individuals. For long-term monitoring, consideration of seasonality, weather, social interactions, and age (Ables, 1969; Dorning and Harris, 2019; Kämmerle et al., 2020; Storm, 1965) in the modeling approach may also be necessary.

Despite initial concerns, the battery life of the implants was demonstrated sufficient for the whole duration of the study and beyond in fox 0fb0. In this work, the recording gap did not compromise the most critical objective of detecting changes early via alarms before any clinical signs were detected. However, for the two foxes that died before data collection resumed (0faf and 0fb1) in particular, the recording gap limited our ability to track the effects of disease development on body temperature and activity. While we did obtain 6-8 consecutive alarms for these two foxes before data collection stopped, it is impossible to know whether or not these alarms would have continued consecutively until death, therefore leaving some uncertainty regarding the true precocity of our approach. In the future, the recording by the implants should be continuous for completeness of data and analyses.

There is growing interest in monitoring individual animals using sensors to track both health and welfare, especially for farm animals (Chapa et al., 2020; van Dixhoorn et al., 2024). This approach can provide early warnings for a range of issues ranging from emerging (Martínez-Avilés et al., 2017; Peña-Mosca et al., 2025) or endemic disease (Dittrich, 2022; Wottlin et al., 2021) to stress (Lardy et al., 2023; Wagner et al., 2021) or lameness (Miekley, 2012; Pastell, 2009; van Hertem, 2013). Regardless of the target issue for early detection, one of the principal challenges of these approaches is validating results and in particular, differentiating between true and false alarms (Garrido et al., 2023). Experimental studies such as ours offer a context for which inoculation dates and incubating periods are generally known and can thus play an important role in the development and validation of such systems, as well as improving the health and welfare of animals, especially uncommon models, used in experimental protocols.

## Acknowledgements

We thank Dr. Delphine Bouard for the placement of the implants and comments on the manuscript. Emka TECHNOLOGIES, France, provided us with a complete digital telemetry system allowing physiological measurements in 4 foxes.

*For the purpose of Open Access, a CC-BY public copyright licence has been applied by the authors to the present document and will be applied to all subsequent versions up to the Author Accepted Manuscript arising from this submission.*

## Data, scripts, code, and supplementary information availability

All data and scripts are available online: doi.org/10.5281/zenodo.18258116 (Shapiro, 2026) and https://github.com/jtshapiro/FoxMonitoring_Temperature_Activity; Supplementary information is available online: https://doi.org/10.5281/zenodo.18267755 (Shapiro et al., 2006).

