## Supplementary Information for "Potential for real-time health and welfare monitoring in experimental rabies infection in red fox (Vulpes vulpes) using implants"

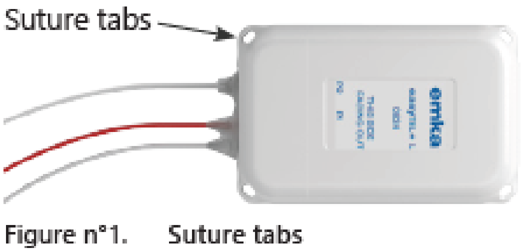


**Figure S1.** An easyTEL+ implant (model ET-M2-ETA - Emka Technologies, France) used in this study.

**Table S1.** Ranking of models (Shewhart and EWMA with varying confidence levels or σ, respectively) for anomaly detection for fox 0fae. Models are ordered by rank starting from the best, based on the lowest number of alarms during the calibration period (Calibration alarms), the greatest number of consecutive alarms before death, and the lowest number of single alarms in the monitoring period. Total alarms indicate total number of alarms during the monitoring period (considered as true positives).

| Measure | Time period | Algorithm | CL / nσ | Calibration alarms | Consecutive alarms | Singleton alarms | Total alarms |
|---|---|---|---|---|---|---|---|
| Activity | Daily | EWMA | 2 | 0 | 10 | 0 | 15 |
| Activity | Day-time | EWMA | 2 | 0 | 9 | 1 | 13 |
| Activity | Daily | EWMA | 3 | 0 | 7 | 1 | 8 |
| Activity | Night-time | EWMA | 2 | 0 | 6 | 0 | 6 |
| Activity | Day-time | EWMA | 3 | 0 | 6 | 1 | 7 |
| Activity | Daily | EWMA | 4 | 0 | 5 | 1 | 6 |
| Activity | Night-time | EWMA | 3 | 0 | 5 | 0 | 5 |
| Activity | Night-time | EWMA | 4 | 0 | 5 | 0 | 5 |
| Activity | Daily | Shewhart | 0.997 | 0 | 5 | 0 | 5 |
| Activity | Daily | Shewhart | 0.9999 | 0 | 5 | 0 | 5 |
| Activity | Day-time | Shewhart | 0.9999 | 0 | 4 | 0 | 4 |
| Activity | Day-time | EWMA | 4 | 0 | 4 | 1 | 5 |
| Activity | Day-time | Shewhart | 0.997 | 0 | 4 | 1 | 5 |
| Activity | Night-time | Shewhart | 0.9999 | 0 | 2 | 0 | 2 |
| Activity | Night-time | Shewhart | 0.997 | 0 | 2 | 1 | 3 |
| Activity | Night-time | Shewhart | 0.95 | 1 | 6 | 0 | 6 |
| Activity | Day-time | Shewhart | 0.95 | 1 | 4 | 3 | 9 |
| Activity | Daily | Shewhart | 0.95 | 2 | 7 | 3 | 10 |
| Temperature | Night-time | EWMA | 4 | 3 | 0 | 0 | 13 |
| Temperature | Night-time | Shewhart | 0.9999 | 2 | 0 | 2 | 9 |
| Temperature | Daily | EWMA | 4 | 4 | 4 | 0 | 14 |
| Temperature | Daily | Shewhart | 0.9999 | 4 | 4 | 2 | 11 |
| Temperature | Night-time | EWMA | 3 | 4 | 0 | 0 | 14 |
| Temperature | Night-time | Shewhart | 0.997 | 4 | 0 | 2 | 9 |
| Temperature | Day-time | EWMA | 4 | 5 | 4 | 0 | 13 |
| Temperature | Day-time | EWMA | 3 | 6 | 5 | 0 | 15 |

| | | | | | | | |
|---|---|---|---|---|---|---|---|
| Temperature | Daily | Shewhart | 0.997 | 6 | 4 | 2 | 13 |
| Temperature | Day-time | Shewhart | 0.9999 | 6 | 4 | 4 | 10 |
| Temperature | Night-time | EWMA | 2 | 6 | 0 | 0 | 15 |
| Temperature | Night-time | Shewhart | 0.95 | 7 | 0 | 3 | 13 |
| Temperature | Day-time | Shewhart | 0.95 | 8 | 7 | 2 | 14 |
| Temperature | Day-time | Shewhart | 0.997 | 8 | 6 | 2 | 13 |
| Temperature | Day-time | EWMA | 2 | 8 | 5 | 0 | 15 |
| Temperature | Daily | EWMA | 2 | 8 | 4 | 0 | 15 |
| Temperature | Daily | EWMA | 3 | 8 | 4 | 0 | 15 |
| Temperature | Daily | Shewhart | 0.95 | 8 | 4 | 1 | 14 |

**Table S2.** Ranking of models (Shewhart and EWMA with varying confidence levels or nσ, respectively) for anomaly detection for fox 0faf. Models are ordered by rank starting from the best, based on the lowest number of alarms during the calibration period (Calibration alarms), the greatest number of consecutive alarms before the end of data collection (8 days before death), and the lowest number of single alarms in the monitoring period. Total alarms indicate total number of alarms during the monitoring period (considered as true positives). Only models that produced at least one alarm are included.

| Measure | Time period | Algorithm | CL /n σ | Calibration alarms | Consecutive alarms | Singleton alarms | Total alarms |
|---|---|---|---|---|---|---|---|
| Activity | Daily | EWMA | 3 | 0 | 6 | 0 | 6 |
| Activity | Daily | EWMA | 4 | 0 | 2 | 2 | 4 |
| Activity | Day-time | EWMA | 2 | 0 | 2 | 0 | 5 |
| Activity | Night-time | EWMA | 4 | 0 | 2 | 0 | 2 |
| Activity | Daily | EWMA | 2 | 1 | 8 | 0 | 8 |
| Temperature | Night-time | Shewhart | 0.9999 | 1 | 0 | 2 | 2 |
| Temperature | Day-time | Shewhart | 0.9999 | 2 | 0 | 1 | 5 |
| Activity | Night-time | EWMA | 3 | 3 | 3 | 0 | 3 |
| Temperature | Daily | Shewhart | 0.997 | 3 | 0 | 1 | 5 |
| Temperature | Daily | Shewhart | 0.9999 | 3 | 0 | 1 | 5 |
| Temperature | Day-time | Shewhart | 0.997 | 3 | 0 | 1 | 5 |
| Temperature | Night-time | Shewhart | 0.997 | 3 | 0 | 2 | 2 |
| Temperature | Day-time | Shewhart | 0.95 | 4 | 0 | 1 | 6 |
| Activity | Night-time | EWMA | 2 | 5 | 4 | 0 | 6 |
| Temperature | Daily | Shewhart | 0.95 | 5 | 0 | 1 | 6 |
| Temperature | Night-time | Shewhart | 0.95 | 5 | 0 | 2 | 4 |

**Table S3.** Ranking of models (Shewhart and EWMA with varying confidence levels or nσ, respectively) for anomaly detection for fox 0fb1. Models are ordered by rank starting from the best, based on the lowest number of alarms during the calibration period (Calibration alarms), the greatest number of consecutive alarms before the end of data collection (12 days before death), and the lowest number of single alarms in the monitoring period. Total alarms indicate total number of alarms during the monitoring period (considered as true positives).

| Measure | Time period | Algorithm | CL / nσ | Consecutive alarms | Calibration alarms | Singleton alarms | Total alarms |
|---|---|---|---|---|---|---|---|
| Activity | Night-time | EWMA | 4 | 8 | 0 | 0 | 8 |
| Activity | Day-time | EWMA | 3 | 3 | 0 | 0 | 5 |
| Temperature | Daily | EWMA | 4 | 5 | 0 | 1 | 6 |
| Temperature | Daily | Shewhart | 0.9999 | 3 | 0 | 3 | 6 |
| Temperature | Day-time | EWMA | 3 | 3 | 0 | 1 | 6 |
| Temperature | Day-time | EWMA | 4 | 3 | 0 | 1 | 4 |
| Activity | Daily | EWMA | 3 | 0 | 0 | 1 | 7 |
| Activity | Daily | EWMA | 4 | 0 | 0 | 0 | 2 |
| Activity | Daily | Shewhart | 0.997 | 0 | 0 | 2 | 2 |
| Activity | Daily | Shewhart | 0.9999 | 0 | 0 | 1 | 1 |
| Activity | Day-time | EWMA | 4 | 0 | 0 | 2 | 2 |
| Activity | Day-time | Shewhart | 0.9999 | 0 | 0 | 2 | 2 |
| Activity | Day-time | EWMA | 2 | 6 | 1 | 0 | 6 |
| Temperature | Daily | EWMA | 3 | 5 | 1 | 0 | 7 |
| Activity | Day-time | Shewhart | 0.997 | 0 | 1 | 2 | 2 |
| Activity | Night-time | Shewhart | 0.9999 | 0 | 1 | 2 | 4 |
| Temperature | Night-time | EWMA | 4 | 3 | 2 | 0 | 3 |
| Activity | Night-time | Shewhart | 0.997 | 0 | 2 | 2 | 6 |
| Temperature | Day-time | Shewhart | 0.9999 | 0 | 2 | 4 | 4 |
| Temperature | Night-time | Shewhart | 0.997 | 0 | 2 | 4 | 4 |
| Temperature | Night-time | Shewhart | 0.9999 | 0 | 2 | 3 | 3 |
| Activity | Night-time | EWMA | 2 | 8 | 3 | 0 | 8 |
| Activity | Night-time | EWMA | 3 | 8 | 3 | 0 | 8 |
| Temperature | Night-time | EWMA | 3 | 6 | 3 | 0 | 6 |
| Temperature | Daily | EWMA | 2 | 5 | 3 | 0 | 7 |

| | | | | | | | |
|---|---|---|---|---|---|---|---|
| Activity | Daily | EWMA | 2 | 0 | 3 | 1 | 8 |
| Activity | Daily | Shewhart | 0.95 | 0 | 3 | 2 | 4 |
| Activity | Day-time | Shewhart | 0.95 | 0 | 3 | 1 | 3 |
| Activity | Night-time | Shewhart | 0.95 | 0 | 3 | 1 | 7 |
| Temperature | Day-time | Shewhart | 0.997 | 0 | 3 | 5 | 5 |
| Temperature | Daily | Shewhart | 0.95 | 3 | 4 | 3 | 6 |
| Temperature | Daily | Shewhart | 0.997 | 3 | 4 | 3 | 6 |
| Temperature | Day-time | EWMA | 2 | 3 | 4 | 1 | 6 |
| Temperature | Day-time | Shewhart | 0.95 | 3 | 5 | 3 | 6 |
| Temperature | Night-time | EWMA | 2 | 6 | 6 | 1 | 7 |
| Temperature | Night-time | Shewhart | 0.95 | 0 | 7 | 3 | 5 |

**Table S4.** Ranking of models (Shewhart and EWMA with varying confidence levels or nσ, respectively) for anomaly detection for fox 0fb0. Models are ordered by rank starting from the best, based on the lowest number of alarms during the calibration period (Calibration alarms) and the lowest number of singleton alarms in the monitoring period. This fox survived inoculation. Total alarms indicate total number of alarms during the monitoring period.

| Measure | Time period | Algorithm | CL / nσ | Calibration alarms | Singleton alarms | Total alarms |
|---|---|---|---|---|---|---|
| Activity | Day-time | EWMA | 4 | 0 | 0 | 21 |
| Activity | Daily | EWMA | 2 | 0 | 1 | 45 |
| Activity | Daily | EWMA | 3 | 0 | 1 | 34 |
| Activity | Day-time | EWMA | 3 | 0 | 1 | 25 |
| Activity | Night-time | EWMA | 2 | 0 | 1 | 39 |
| Activity | Night-time | EWMA | 3 | 0 | 1 | 31 |
| Temperature | Daily | EWMA | 4 | 0 | 1 | 49 |
| Activity | Daily | EWMA | 4 | 0 | 2 | 25 |
| Activity | Daily | Shewhart | 0.997 | 0 | 2 | 16 |
| Activity | Day-time | EWMA | 2 | 0 | 2 | 30 |
| Activity | Day-time | Shewhart | 0.9999 | 0 | 2 | 14 |
| Activity | Night-time | EWMA | 4 | 0 | 2 | 23 |
| Activity | Night-time | Shewhart | 0.9999 | 0 | 2 | 2 |
| Activity | Daily | Shewhart | 0.95 | 0 | 3 | 29 |
| Activity | Night-time | Shewhart | 0.997 | 0 | 3 | 9 |
| Activity | Daily | Shewhart | 0.9999 | 0 | 4 | 9 |
| Activity | Day-time | Shewhart | 0.997 | 0 | 5 | 21 |
| Activity | Night-time | Shewhart | 0.95 | 0 | 6 | 26 |
| Activity | Day-time | Shewhart | 0.95 | 1 | 2 | 27 |
| Temperature | Daily | Shewhart | 0.9999 | 1 | 6 | 30 |
| Temperature | Daily | EWMA | 3 | 2 | 2 | 54 |
| Temperature | Day-time | EWMA | 4 | 2 | 2 | 44 |
| Temperature | Night-time | Shewhart | 0.9999 | 2 | 4 | 21 |
| Temperature | Daily | Shewhart | 0.997 | 3 | 2 | 40 |
| Temperature | Day-time | EWMA | 3 | 3 | 2 | 49 |
| Temperature | Day-time | Shewhart | 0.9999 | 3 | 6 | 29 |

| | | | | | | |
|---|---|---|---|---|---|---|
| Temperature | Day-time | Shewhart | 0.997 | 4 | 2 | 39 |
| Temperature | Daily | EWMA | 2 | 5 | 1 | 57 |
| Temperature | Daily | Shewhart | 0.95 | 6 | 2 | 49 |
| Temperature | Day-time | Shewhart | 0.95 | 6 | 2 | 49 |
| Temperature | Night-time | EWMA | 4 | 6 | 6 | 26 |
| Temperature | Night-time | Shewhart | 0.997 | 6 | 7 | 26 |
| Temperature | Day-time | EWMA | 2 | 7 | 1 | 53 |
| Temperature | Night-time | EWMA | 3 | 7 | 4 | 32 |
| Temperature | Night-time | Shewhart | 0.95 | 7 | 7 | 33 |
| Temperature | Night-time | EWMA | 2 | 8 | 4 | 40 |
